# The Protein Interaction Networks of Catalytically-Active and Catalytically-Inactive PqsE in *Pseudomonas aeruginosa*

**DOI:** 10.1101/2022.05.31.494267

**Authors:** Isabelle R. Taylor, Laura A. Murray-Nerger, Ileana M. Cristea, Bonnie L. Bassler

## Abstract

*Pseudomonas aeruginosa* is a human pathogen that relies on quorum sensing to establish infections. The PqsE quorum-sensing protein is required for *P. aeruginosa* virulence factor production and infection. PqsE has a reported enzymatic function in the biosynthesis of the quorum-sensing autoinducer called PQS. However, this activity is redundant because, in the absence of PqsE, this role is fulfilled by alternative thioesterases. Rather, PqsE drives *P. aeruginosa* pathogenic traits via a protein-protein interaction with the quorum-sensing receptor/transcription factor, RhlR, an interaction that enhances affinity of RhlR for target DNA sequences. PqsE catalytic activity is dispensable for interaction with RhlR. Thus, PqsE virulence function can be decoupled from its catalytic function. Here, we present an immunoprecipitation-mass spectrometry method employing eGFP-PqsE fusions to define the protein interactomes of wildtype PqsE and the catalytically inactive PqsE(D73A) variant in *P. aeruginosa* and their dependence on RhlR. Several proteins were identified to have specific interactions with wildtype PqsE, while not forming associations with PqsE(D73A). In the Δ*rhlR* strain, an increased number of specific PqsE interactors were identified, including the partner autoinducer synthase to RhlR, called RhlI. Collectively, these results suggest that specific protein-protein interactions depend on PqsE catalytic activity and that RhlR may prevent proteins from interacting with PqsE, possibly due to competition between RhlR and other proteins for PqsE binding. Our results provide a foundation for the identification of the *in vivo* PqsE catalytic function and, potentially, new proteins involved in *P. aeruginosa* quorum sensing.

**IMPORTANCE:** *Pseudomonas aeruginosa* causes hospital-borne infections in vulnerable patients, including in immunocompromised individuals, burn victims, and cancer patients undergoing chemotherapy. There are no effective treatments for *P. aeruginosa* infections, which are usually broadly resistant to antibiotics. Animal models show that to establish infection and to cause illness, *P. aeruginosa* relies on an interaction between two proteins: PqsE and RhlR. There could be additional protein-protein interactions involving PqsE, which, if defined, could be exploited for the design of new therapeutic strategies to combat *P. aeruginosa*. Here, we reveal previously unknown protein interactions in which PqsE participates that will be investigated for potential roles in pathogenesis.

## OBSERVATION

The opportunistic human pathogen *Pseudomonas aeruginosa* is responsible for causing highly antibiotic resistant, virtually untreatable nosocomial infections (1, 2). *P. aeruginosa* pathogenic traits such as virulence factor production and biofilm formation are under control of the bacterial cell-to-cell communication process called quorum sensing (QS) (3). QS relies on the production, release, and group-wide detection of signal molecules called autoinducers. The QS network in *P. aeruginosa* is composed of multiple, interconnecting branches, including two acyl homoserine lactone autoinducer synthase/receptor-transcription factor pairs, LasI/LasR and RhlI/RhlR (4). The RhlI/RhlR pair, responsible for, respectively, producing and detecting the C4-homoserine lactone autoinducer (C4-HSL), controls cell density-dependent gene expression (5, 6). Curiously, given that RhlR and RhlI function in a receptor-ligand partnership, pathogenic phenotypes resulting from deletion of *rhlR* differ from those following deletion of *rhlI*. For instance, a Δ*rhlI* mutant can mount an infection in a murine host, whereas a Δ*rhlR* mutant is avirulent and does not establish infection (7). Deletion of the gene encoding the enzyme, PqsE, that acts in a different *P. aeruginosa* QS pathway, results in the identical loss of pathogenic phenotypes as occur following deletion of *rhlR* (8). Consistent with this result, we recently showed that PqsE and RhlR make a protein-protein interaction that enhances the affinity of RhlR for target DNA sequences (9, 10). The PqsE-RhlR interaction is, moreover, essential for RhlR-controlled QS phenotypes. Furthermore, the PqsE-RhlR interaction does not depend on PqsE catalytic activity, as the catalytically inactive PqsE(D73A) variant is capable of interacting with RhlR *in vitro* and driving virulence phenotypes in *P. aeruginosa* (9).

The *in vivo* role of PqsE catalytic function remains unknown. PqsE is encoded by the fifth gene in the *pqsABCDE* operon (11). PqsA-E together with PqsH (encoded separately in the genome) are responsible for synthesis of the *Pseudomonas* Quinolone Signal (PQS), a QS autoinducer (11, 12). However, a Δ*pqsE* mutant produces wildtype levels of PQS, and alternative thioesterases have been shown to fulfill the reported biosynthetic function of PqsE in this pathway (the conversion of 2-aminobenzoyl acetyl-CoA to 2-aminobenzoyl acetate) (13). Thus, beyond PqsE catalytic function being unnecessary for interaction with RhlR, it is also dispensable for production of PQS, highlighting the possibility that there exist as-of-yet unknown roles for PqsE-driven catalysis.

In this work, we explore the question of whether the PqsE-RhlR interaction occurs *in vivo* and whether PqsE participates in additional protein-protein interactions. To probe these possibilities, we determine the PqsE interactome in *P. aeruginosa* PA14. Another goal of this study is to gain insight into possible biosynthetic pathways requiring PqsE catalytic function. Here, we develop an immunoaffinity purification-mass spectrometry (IP-MS) strategy using PqsE tagged with monomeric eGFP as the bait protein. We employ both wildtype PqsE (designated PqsE(WT)) and a catalytically dead version harboring the D73A mutation (designated PqsE(D73A)) to distinguish PqsE interactions that require intact catalytic function from those that do not. The PqsE(WT) and PqsE(D73A) interactomes are identified in *P. aeruginosa* PA14 and in a Δ*rhlR* mutant strain. The results reveal a set of PqsE-protein interactions that depend on intact PqsE catalytic function. Furthermore, the Δ*rhlR* strain analyses show that the PqsE-RhlR interaction may prevent PqsE from interacting with a diverse set of proteins, including RhlI. These findings provide a platform to begin to address the as-of-yet unknown role(s) PqsE catalysis plays *in vivo* and to identify new proteins and mechanisms involved in the *P. aeruginosa* QS network.

### DESIGNING THE PQSE IP-MS WORKFLOW

The PqsE-RhlR interaction is essential for *P. aeruginosa* to produce the toxin pyocyanin (9, 10). Thus, we monitored pyocyanin production to guide our design of a functional affinity tagged PqsE fusion as the bait protein for IP-MS experiments to define the PqsE interactome. Both N-terminal (eGFP-PqsE) and C-terminal (PqsE-eGFP) tagged constructs were engineered into the pUCP18 vector and expressed in *ΔpqsE P. aeruginosa* PA14. Pyocyanin production was measured and compared to that from a strain carrying untagged PqsE on the same plasmid (Figure S1a). The C-terminally tagged PqsE-eGFP construct drove significantly reduced pyocyanin production (22% compared to untagged), whereas the N-terminally tagged eGFP-PqsE showed only a small reduction in pyocyanin production (83% compared to untagged). This result is consistent with the finding that substitution of residues near the C-terminus of PqsE (R243A/R246A/R247A) abolishes interaction with RhlR (10). Therefore, we used eGFP-PqsE constructs as bait in our IP-MS experiments. The eGFP tag in each construct was also confirmed to be folded and functional as judged by fluorescence output (Figure S1b). To control for interactions involving the eGFP tag, eGFP alone was also cloned into pUCP18 and identically assayed by IP-MS. In analyzing the data from each IP-MS experiment, proteins were considered specific PqsE-interactors if they were enriched by at least two-fold compared to their abundance in the eGFP alone sample among other cut-off criteria that are described in the Methods. We know that PqsE interacts with RhlR, so we performed this set of experiments in both WT and Δ*rhlR P. aeruginosa* strains to examine whether the presence/absence of RhlR influenced which specific interactions occur with PqsE. All strains used in this study are described in Table S1.

### PQSE PROTEIN INTERACTIONS THAT DEPEND ON CATALYTIC FUNCTION

In wildtype *P. aeruginosa* PA14, 11 proteins were identified as enriched in the eGFP-PqsE(WT) IP-MS experiment compared to that with eGFP alone (Figure 1, Table S2). This result represents 0.2% of the annotated *P. aeruginosa* proteome, and therefore indicated highly specific interactions. Indeed, RhlR was identified as an interacting partner for PqsE. RhlR was also identified in the eGFP-PqsE(D73A) IP-MS experiment, confirming that this interaction is independent of the PqsE catalytic function. In contrast, seven of the 11 proteins identified in the eGFP-PqsE(WT) experiment did not pass specificity filtering in the eGFP-PqsE(D73A) IP-MS analysis (ovals, Figure 1), suggesting that their interactions with PqsE depend on PqsE catalytic function. These seven proteins are UreA, ThrB, GpsA, AcpP, ApeB, PA14_14020, and YbeY. Of these potential interacting partners, the acyl carrier protein AcpP is of particular interest as it participates in the synthesis of acyl homoserine lactone QS autoinducers (14). More generally, all of the proteins whose interactions with PqsE depended on it possessing intact catalytic function are enzymes, suggesting that their interactions with PqsE could potentially accomplish a biosynthetic function. In addition to RhlR, the proteins that interacted with both PqsE(WT) and the catalytically dead PqsE(D73A) protein were GroL, GcvH1, and MaiA. The molecular chaperone, DnaK, was also observed as a specific interactor with PqsE(D73A), but not with PqsE(WT). As GroL and DnaK are both chaperones, highly abundant, and promiscuous interactors, their interactions with PqsE are to be considered cautiously.

**Figure 1:**
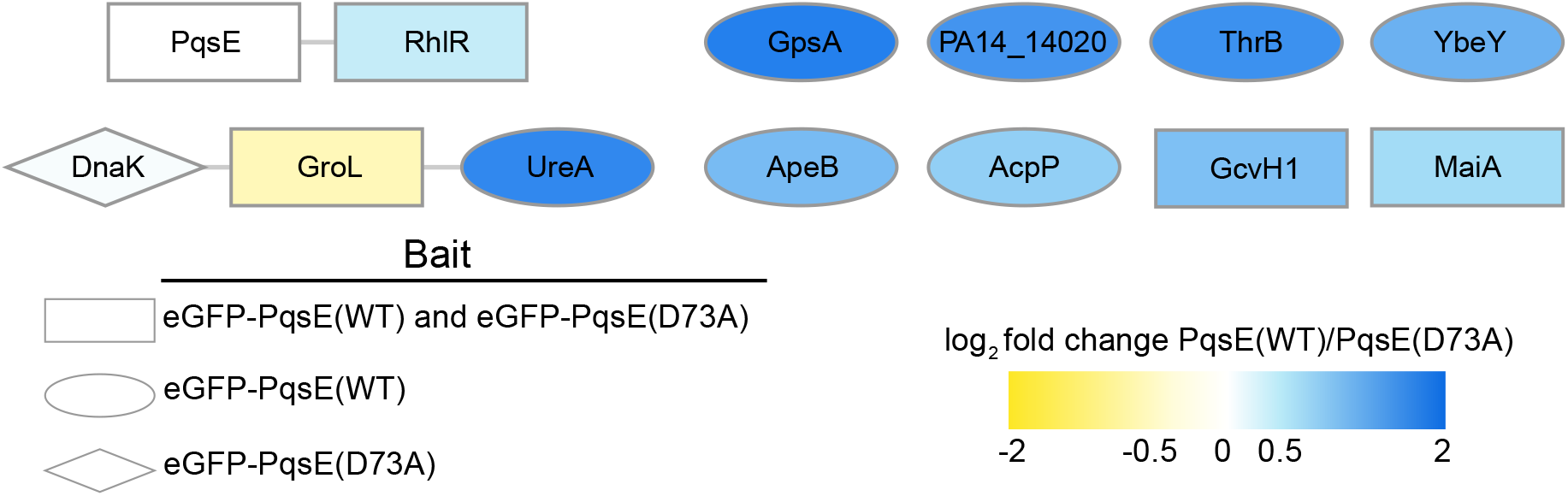
Specificity-filtered interactions with either eGFP-PqsE(WT) or eGFP-PqsE(D73A) in wildtype *P. aeruginosa* PA14. Shapes indicate whether the interaction passed specificity filtering for both bait proteins (rectangles), only eGFP-PqsE(WT) (ovals), or only eGFP-PqsE(D73A) (diamonds). Gray lines represent known or predicted interactions from the STRING database. Nodes are colored by abundance-based enrichment in the eGFP-PqsE(WT) (blue) or eGFP-PqsE(D73A) (yellow) IP experiments.

### RHLR INHIBITS OTHER PROTEINS FROM INTERACTING WITH PQSE

When the above IP-MS experiment was conducted with eGFP-PqsE(WT) in *ΔrhlR P. aeruginosa* PA14, surprisingly, the number of PqsE-specific interacting proteins increased to 36 (Figure 2; see rectangles and ovals, Table S3), 13 of which were at least two-fold more abundant in the Δ*rhlR* strain than in wildtype *P. aeruginosa* PA14 (Figure S2a; see blue nodes, Table S4). With the exceptions of GroL and DnaK, none of the proteins that passed specificity filtering in wildtype *P. aeruginosa* PA14 were identified as specific PqsE interactors in the Δ*rhlR* strain. We note particularly that, in *ΔrhlR P. aeruginosa* PA14, both eGFP-PqsE(WT) and eGFP-PqsE(D73A) interacted with the C4-HSL synthase, RhlI (Figure 2, Figure S2a,b, Tables S4 and S5). This finding suggests that the potential PqsE-RhlI interaction is independent of PqsE catalytic function. In contrast to RhlI, several of the eGFP-PqsE(WT) interactors identified in the Δ*rhlR* strain did not pass filtering in the Δ*rhlR* strain with eGFP-PqsE(D73A) as the bait (Figure 2; ovals), suggesting that these interactions depend on PqsE catalytic function.

**Figure 2:**
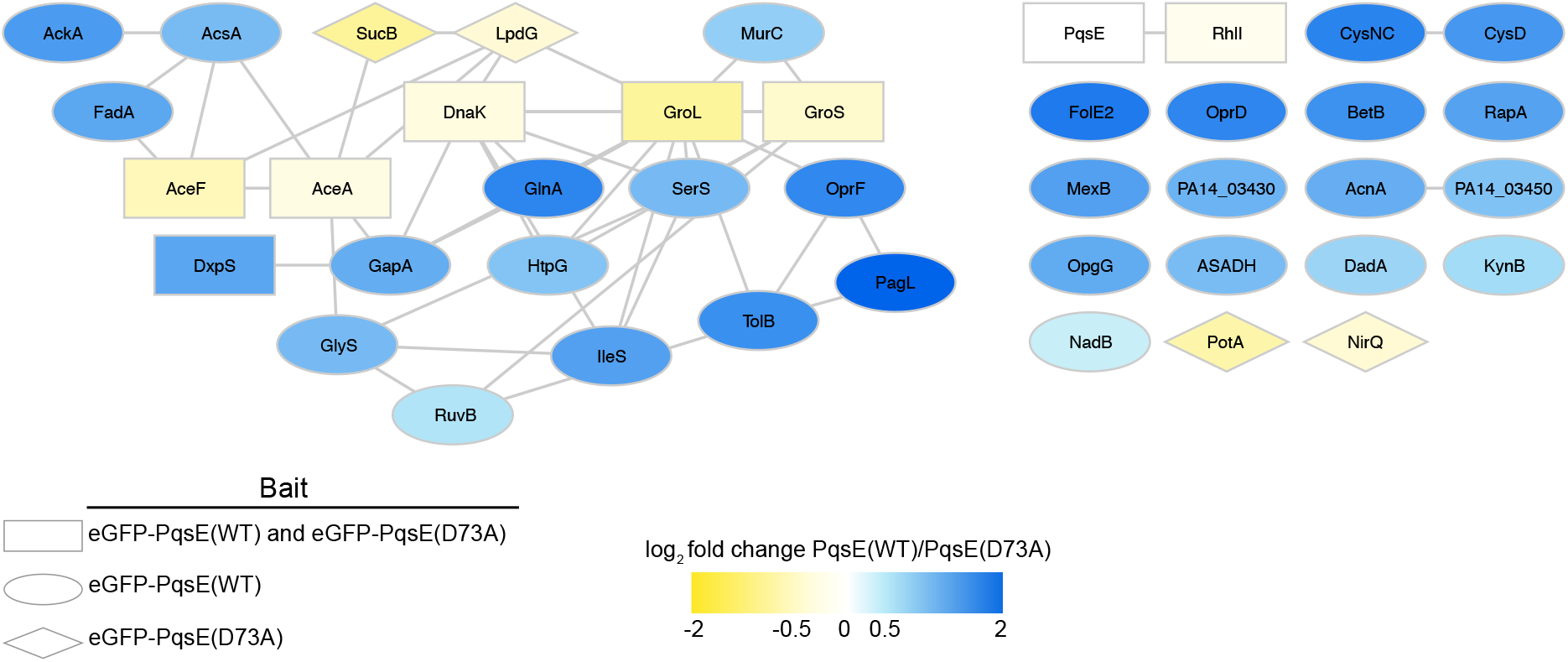
Specificity-filtered interactions with eGFP-PqsE(WT) or eGFP-PqsE(D73A) in the *P. aeruginosa* PA14 Δ*rhlR* strain. Shapes indicate whether the interaction passed specificity filtering for both bait proteins (rectangles), only eGFP-PqsE(WT) (ovals), or only eGFP-PqsE(D73A) (diamonds). Gray lines represent known or predicted interactions from the STRING database. Nodes are colored by abundance-based enrichment in the eGFP-PqsE(WT) (blue) or eGFP-PqsE(D73A) (yellow) IP experiments.

## DISCUSSION

PqsE plays an essential role in driving pathogenic behaviors in *P. aeruginosa* PA14, highlighting the importance of defining its *in vivo* function. We know that to activate virulence, PqsE makes a protein-protein interaction with the QS regulator, RhlR, and this interaction does not rely on PqsE catalytic activity. The PqsE *in vivo* catalytic function, as well as any additional protein-protein interactions, remain unknown. Here, we engineered a functional (i.e., capable of driving pyocyanin production, Figure S1a) PqsE protein with eGFP fused to the N-terminus for use in the *in vivo* IP-MS experiments. Future experiments could employ PqsE constructs with eGFP fused to the C-terminus to identify interactions involving the N-terminus of PqsE. Through our IP-MS analyses, we have identified additional PqsE interactions that occur in *P. aeruginosa* PA14, and we can distinguish those that require intact PqsE catalytic function from those that do not. Proteins that interact with PqsE(WT), but not the catalytically dead PqsE(D73A) variant, could be indicative of biosynthetic pathways in which PqsE participates. Notably, the AcpP acyl carrier protein was identified, as was the RhlI C4-HSL synthase in the Δ*rhlR* dataset. These findings hint that the annotated thioesterase function of PqsE could be important for editing acyl chain length during the synthesis of acyl homoserine lactone autoinducers, such as C4-HSL. We are currently pursuing metabolomic analyses to identify small molecule substrates and products of PqsE.

The interaction between PqsE and RhlR was previously established through mutagenesis of PqsE combined with *in vitro* pull-down assays using recombinant proteins produced in *Escherichia coli* (9, 10). Here, we validate that the PqsE-RhlR interaction occurs in *P. aeruginosa* PA14 *in vivo*, and furthermore, that it is independent of PqsE catalytic function. Surprisingly, we also show that PqsE interacts with many more proteins in the Δ*rhlR* strain than in wildtype *P. aeruginosa* PA14. These putative PqsE interactors harbor a wide diversity of functions. This result suggests that interaction between PqsE and RhlR is primary and it blocks interactions that PqsE can undertake with other proteins, including RhlI. Perhaps RhlI and RhlR compete for interaction with PqsE, and their relative production levels under different conditions determine which complex will form. Although these PqsE interactions remain to be validated, the results of this study provide the starting point for exploring the *in vivo* functions of this vital component of the *P. aeruginosa* QS and pathogenesis networks.

## METHODS

### GROWTH CONDITIONS AND SAMPLE PREPARATION

We use the designation PA14 to signify *P. aeruginosa* UCBPP-PA14. All strains used in this study are listed in the Supplementary Table S1. The following six strains were grown as overnight cultures in LB supplemented with carbenicillin (400 µg/mL): WT PA14 + pUCP18_eGFP, WT PA14 + pUCP18_eGFP-PqsE(WT), WT PA14 + pUCP18_eGFP-PqsE(D73A), Δ*rhlR* PA14 + pUCP18_eGFP, Δ*rhlR* PA14 + pUCP18_eGFP-PqsE(WT), and Δ*rhlR* PA14 + pUCP18_eGFP-PqsE(D73A). The overnight cultures were back-diluted 1:100 into 25 mL fresh LB + carbenicillin and grown at 37 °C with shaking (200 rpm) until they reached OD_600_ = 1.5. Cells were pelleted by centrifugation at 4,000 rpm for 15 min, washed three times with 5 mL PBS, resuspended in 200 µL freezing buffer (20 mM HEPES, 1.2% polyvinylpyrrolidone (w/v), pH 7.4), and flash-frozen by slowly pipetting droplets into liquid nitrogen. The frozen droplets were stored at -80 °C until cryogenic grinding. Cryogenic grinding was performed in a Retsch CryoMill with nine cycles at a frequency of 30 Hz lasting 1.5 min per cycle. After grinding, the frozen cell powders were transferred into pre-chilled LoBind tubes (Amuza, Inc. Eicom, USA) and stored at -80 °C until lysis and immunoaffinity purification were performed. All samples were collected in biological triplicate.

### LYSIS AND IMMUNOAFFINITY PURIFICATION (IP)

The frozen cell powders were resuspended in pre-chilled lysis buffer (20 mM HEPES pH 7.4, 100 mM potassium acetate, 2 mM MgCl_2_, 0.1% Tween-20 (v/v), 1 µM ZnCl_2_, 1 µM CaCl_2_, 1% Triton-X100 (v/v), 200 mM NaCl, 0.5 mM phenylmethylsulfonyl fluoride (PMSF), 1:2,500 Pierce Universal Nuclease, and a protease inhibitor cocktail (Roche, one tablet/10 mL buffer)). Resuspension was carried out by inversion and gentle vortex, followed by rotation for 30 min at 4 °C until samples were completely solubilized. Lysates were subsequently subjected to Polytron homogenization by pulsing twice for 15 s at a speed of 22,500 rpm. Samples were incubated on ice for 10 s between pulses. Lysates were cleared by centrifugation at 10,000 *x g* at 4 °C for 10 min. Protein concentration was determined by bicinchoninic acid analysis, and 500 µg of protein from each sample was incubated with 35 µL pre-washed GFP-Trap magnetic beads (Chromotek, Inc.) for 1 h at 4 °C with rotation (1 mg/mL final lysate concentration). The beads were separated from the total sample using a magnet, and supernatant was removed by aspiration. The beads were washed three times with 500 µL wash buffer (lysis buffer lacking PMSF, nuclease, and protease inhibitors). A final wash with cold PBS was performed, and the beads were transferred to a new tube. Proteins were eluted in 50 µL 1x TES buffer (2% SDS, 0.5 mM EDTA, 53 mM Tris HCl, 70 mM Tris Base) by incubation at 70 °C for 10 min and vortex for 20 s. The eluate was transferred to a new Lo-Bind tube.

### PREPARATION OF IP SAMPLES FOR MASS SPECTROMETRY ANALYSIS

IP eluates were concentrated two-fold in a speedvac. Proteins were reduced and alkylated with 25 mM Tris(2-carboxyethyl)phosphine (TCEP) (Thermo Fisher) and 50 mM chloroacetamide (CAM) (Fisher Scientific) by heating for 20 min at 70 °C. The resulting samples were digested using an S-Trap micro column (Protifi LLC). Briefly, samples were acidified to 1.2% phosphoric acid, diluted into S-Trap binding buffer (100 mM TEAB, pH 7.1 in 90% methanol), and bound to the S-Trap column by centrifugation at 4,000 x *g* for 30 s. S-Trap-bound sample was washed using the same centrifugation procedure as follows: two washes with S-Trap binding buffer, five washes with methanol/chloroform (4:1 v/v), and three washes with S-Trap binding buffer. Samples were next digested on the S-Trap column with 2.5 µg trypsin diluted in digestion buffer (25 mM TEAB) for 1 h at 47 °C. Trypsinized peptides were eluted through a three-part elution (40 µL 25 mM TEAB, 40 µL 0.2% formic acid, 70 µL 50% acetonitrile/0.2% formic acid) by sequentially adding the elution buffers to the column followed by centrifugation as described above after addition of each buffer. The eluted peptides were pooled in an LC-MS autosampler vial (Fisher Scientific) and dried via speedvac. Peptides were resuspended in 6 µL 1% formic acid (FA)/1% acetonitrile (ACN).

### MASS SPECTROMETRY AND DATA ACQUISITION

IP samples were analyzed on a Q-Exactive HF mass spectrometer (ThermoFisher Scientific) equipped with a Nanospray Flex Ion Source (ThermoFisher Scientific). Peptides were separated on a 50 cm column (360 μm od, 75 μm id, Fisher Scientific) packed in house with ReproSil-Pur C18 (120 Å pore size, 1.9 μm particle size, ESI Source Solutions). Peptides were separated over a 150 min gradient of 3% B to 35% B (solvent A: 0.1% FA, solvent B: 0.1% FA, 97% ACN) at a flow rate of 0.25 nL/min. MS1 scans were collected with the following parameters: 120,000 resolution, 30 ms MIT, 3e6 automatic gain control (AGC), scan range 350 to 1800 m/z, and data collected in profile. MS2 scans were collected with the following parameters: 30,000 resolution, 150 ms MIT, 1e5 AGC, 1.6 m/z isolation window, loop count of 10, NCE of 28, 100.0 m/z fixed first mass, peptide match set to preferred, and data collected in centroid and at a dynamic exclusion of 45 s.

### MASS SPECTROMETRY DATA ANALYSIS

MS/MS spectra were analyzed in Proteome Discoverer v2.4 (Thermo Fisher Scientific). Sequest HT was used to search spectra against a Uniprot database containing *P. aeruginosa* protein sequences (downloaded October 2020) and common contaminants. Offline mass recalibration was performed via the Spectrum Files RC node, and the Minora Feature Detector node was used for label-free MS1 quantification. Fully tryptic peptides with a maximum of two missed cleavages, a 4 ppm precursor mass tolerance, and a 0.02 Da fragment mass tolerance were used in the search. Posttranslational modifications (PTMs) that were allowed included the static modification carbamidomethylation of cysteine and the following dynamic modifications: oxidation of methionine, deamidation of asparagine, loss of methionine plus acetylation of the N-terminus of the protein, acetylation of lysine, and phosphorylation of serine, threonine, and tyrosine. Peptide spectrum match (PSM) validation was accomplished using the Percolator node and PTM sites were assigned in the ptmRS node. PSMs were assembled into peptide and protein identifications with a false discovery rate of less than 1% at both the peptide and protein levels with at least two unique peptides identified per protein. Precursor quantitation required identification in at least two out of the three replicates. Samples were normalized in a retention time dependent manner, imputation was performed using low abundance resampling, protein abundances were calculated using summed abundances, and protein ratio calculations were performed using pairwise ratios.

Data were exported to Microsoft Excel for further processing. In order for a protein to be considered a putative PqsE interactor, the following requirements had to be met: (1) there must be at least a two-fold enrichment of the protein in the sample of interest compared to the eGFP control, (2) at least two peptides had to be quantified in the sample of interest (PqsE(WT) or PqsE(D73A)) in all replicates, and (3) the Grouped Coefficient of Variation (CV%) had to be less than 75%. To compare PqsE(WT) and PqsE(D73A) interactions, bait normalization was performed by dividing the PqsE(WT)/PqsE(D73A) abundance ratio for each interacting protein by the bait abundance ratio for PqsE(WT)/PqsE(D73A). Fold changes of 1.5 or higher were considered significant. Bait normalization was only performed for interactions that passed specificity filtering, as described above. To compare interactions between the wildtype and Δ*rhlR* strains, bait normalization could not be performed, so the wildtype/Δ*rhlR* abundance ratio was calculated, ratios for proteins identified exclusively in one sample were manually verified, and a stringent cutoff of fold changes of 2 or higher were considered significant. Protein interaction networks were generated using STRING (v.11) (15) and Cytoscape (v.3.8.2) (16).

## DATA AVAILABILITY

The mass spectrometry proteomics data have been deposited to the ProteomeXchange Consortium via the PRIDE partner repository with the dataset identifier PXD034149. The reviewer login credentials are as follows:

Username

Password: C5UznktV

## ACKNOWLEDGMENTS

We thank members of the Bassler and Cristea laboratories for helpful advice and discussions. We thank Todd Greco for providing valuable insight. This work was supported by the Howard Hughes Medical Institute, NIH grant 2R37GM065859, and National Science Foundation grant MCB-2043238 to B.L.B., NIH grant F32GM134583 to I.R.T, NIH grant GM114141 to I.M.C., and National Science Foundation grant DGE-1656466 to L.A.M.N. The content herein is solely the responsibility of the authors and does not represent the official views of the National Institutes of Health. The authors declare that they have no competing financial interests.

I.R.T. and L.A.M.N. conducted experiments; I.R.T., L.A.M.N., I.M.C., and B.L.B. designed experiments and prepared the manuscript.

